# Social transmission of inflammation in mice

**DOI:** 10.1101/2024.02.29.582723

**Authors:** Silvia Castany Quintana, Priscila Batista Da Rosa, Kiseko Shionoya, Anders Blomqvist, David Engblom

**Affiliations:** Center for Social and Affective Neuroscience, Department of Biomedical and Clinical Sciences, Linköping University, 58185 Linköping, Sweden; Division of Neurobiology, Department of Biomedical and Clinical Sciences, Linköping University, 58185 Linköping, Sweden

## Abstract

The ability to detect and respond to sickness in others promotes survival. Here we show that mouse dams respond to immune challenged pups by mirroring their inflammatory response. Thus, dams with pups subjected to immune challenge displayed a marked induction of inflammatory mediators in both the brain and the periphery, accompanied by an increase in maternal behaviors and corticosterone levels. This social transmission of inflammation did not require physical contact, and it contributed to the stress hormone response in the dams. In adult dyads, interaction with an immune challenged cagemate did not elicit robust inflammatory signaling but induced an increased responsiveness to a subsequent immune challenge. The identification of social transmission of inflammation, or inflammatory responsiveness, may open new avenues for research on social behavior, just like the description of similar phenomena such as observational fear and transmitted pain have done.

## Introduction

In many species, individuals react to the physiological and affective state of conspecifics. In certain cases, there is social transmission or transfer of the affective state to the interaction partner, leading to a form of affective mirroring. For example, social interaction with somebody who is stressed or in pain can lead to transmission of the stress or the pain-related state to the naïve partner, or “bystander” (1-4). Such transmission occurs both in humans and rodents and might prepare the bystander for coming challenges.

Immune challenge is a strong threat, both for the affected individual and bystanders. For the latter, it could be important to detect the sickness to be able to either avoid or support the affected individual. In these cases, kinship often drives the interaction in the supportive direction (5,6). While it is established that bystanders can detect sickness in others and adapt their behavior accordingly (7,8), it is not known if there is a component of transmission or mirroring of the inflammatory state. Such a transfer or mirroring could potentially prime the immune function of the bystander and possibly also affect the hormonal status and behavior.

## Material and methods

### Animals

C57Bl6J female wild type mice from Janvier Labs (Le Genest-Saint-Isle, France) were used in all experiments. Pregnant females were housed individually in standard individually ventilated cages until pups were born. In all cases, this was the first pregnancy of the dam. Experiments were performed postnatal day (PND) 8–9 when dams were 13-14 weeks old. The number of pups per litter ranged from 4 to 9. Mice were kept under controlled conditions of humidity (40-60 %) and temperature (22±1 °C) on a 12 h/12 h light/dark cycle (lights on at 7 A.M.) with food and water available ad libitum. All animal experiments were approved by the Animal Care and Use Committee at Linköping University.

### Experimental design

Dams with litters were moved to a room with the temperature set at 26–27 °C at least 30 min before the experiment. In a first set of experiments, pups PND 8-9 were given an intraperitoneal (i.p.) injection of lipopolysaccharide (LPS, 40 μg/kg; E. coli O111:B4, Sigma; dissolved in sterile saline) or vehicle (saline). Immediately after injection, the pups were placed back into their home cage and allowed interaction with their dam for 3 h. During that time, maternal behavior was recorded and scored. Subsequently, dams were subjected to a pup retrieval test or sacrificed for molecular and histological analysis.

A second set of experiments was performed in a similar manner to that described above, except that after injections, pups were placed separated from the dam in a new clean cage next to hers. Each cage was covered by a plastic cover with holes. The dam cage was separated from the pup cage by about 5cm. The dam and their pups could see, smell, and hear each other but they could not have physical contact. An additional control group was included where dam and pups were left together and undisturbed (e.g. no injection). After 3 h, dams were sacrificed and samples from brain, liver, and plasma were taken for quantitative PCR, ELISA, and immunohistochemistry. Finally, in a separate set of experiments, pup litters were injected with LPS or saline, as previously described, and dams were injected with indomethacin (10 mg/kg, i.p.; Sigma-Aldrich) or vehicle (NaOH 4% in saline, i.p.), and pups and dams were then allowed to interact for 3 h.

To avoid any possible auditory, visual or olfactory signaling from the sick pups to the control groups, the two experimental conditions (dams exposed to LPS-injected pups and dams exposed to saline-injected pups, respectively) were conducted on different days, with the order of the treatments balanced between the replicates.

To examine social transmission of inflammation between adult dyads, two conspecific female mice were housed together for at least 2 weeks. Female pairs were move to a room with the temperature set at 22±1 °C at least 30 min before the start of the experiment. One of females was injected with 100 μg/kg LPS (or saline) and exposed to the cage-mate for 4 h. After that, the cage-mate was immune challenged by i.p. injection of 10 μg/kg LPS (or injected with saline) and sacrificed 2 h later for molecular analysis.

### Behavioral tests and analysis

Dam and pup interaction was recorded for 3 h after LPS or saline injections. During this time, the standard cage lid was replaced with a transparent cover with holes to allow recording and scoring of maternal care. Behavioral scoring was performed during 40-60 min, 80-100 min, and 120-140 min after injection. The dam behaviors evaluated were pup-directed behaviors (pup-nursing, pup-grooming and time spent in the nest), as well as non-pup directed behaviors, aimed at self-care and maintenance (self-grooming and digging). Nursing behavior was defined as the time that the dam was over the pups, relatively immobile in an arch posture supported by rigid fore- or hind-limbs, with pups attached to the nipples. Pup grooming was defined as the time that the dam spent touching any part of the pup’s body with her tongue, nose or forepaws, or stretching her nose in the direction of the pups. Time spent in the nest was defined as the time the dam spent physically inside the nest in an active or passive behavior. Digging was defined as a series of fast alternating movements of the forepaws scraping back bedding material resulting in accumulation of bedding in a pile under the abdomen of the animal. The behavior was scored as self-grooming when the dam was performing a sequence of cleaning steps by licking and scratching its own fur.

Pup retrieval behavior was tested after 3 h of exposure to pups that had been injected with LPS of saline. Pups were separated from the dams and positioned at the opposite side of the nest. The time to retrieve each pup was scored. Dams that did not retrieve any pup or took more than five minutes to retrieve the first pup were excluded. In the present experiment, 3 out of 15 dams exposed to saline-injected pups and 5 out of 16 dams exposed to LPS-injected pups did not retrieve their pups.

### Quantitative PCR

Dams were sacrificed by asphyxiation with CO_2_ followed by cervical dislocation. The hypothalamus and liver were dissected, immersed in RNAlater (Qiagen; Hilden, Germany) and stored at -20°C for later analysis. RNA was extracted using RNeasy Lipid Tissue Kit (Qiagen) according to the manufacturer’s instructions. The RNA concentrations and quality were measured with a NanoDrop spectrophotometer (ThermoFisher Scientific; <u>Waltham, MA, USA</u>). Only RNA samples with A260/A280 and A260/A230 ratios > 1.8 were included in the experiment. cDNA synthesis was performed with High-Capacity cDNA Reverse Transcription Kit (Applied Biosystems; <u>Waltham, MA, USA</u>). For quantitative PCR, TaqMan Gene Expression Master Mix (Applied Biosystems) was used together with the following TaqMan gene expression assays : Cyclooxygenase-2 (*Ptgs2*;Mm00478374_m1), Interleukin-1β (*Il1b*; Mm01336189_m1), tumor necrosis factor (*Tnf*; Mm00443258_m1) and interleukin-6 (*Il6*; Mm00446190_m1), C-X-C motif chemokine ligand 10 (*Cxcl10*; Mm00445235_m1), and C-C Motif Chemokine Ligand 2 (*Ccl2*; Mm00441242_m1). As endogenous control, Glyceraldehyde 3-phosphate dehydrogenase (*Gapdh*; Mm99999915_g1) was used. Reactions were performed in a Real-Time 7900 Fast apparatus (Applied Biosystems). Relative quantification was carried out using the 2^ΔΔCT^ method. Gene expression changes are expressed as fold change versus the respective control group.

### Corticosterone assay

Trunk-blood was collected in EDTA tubes. Blood samples were centrifuged at 5000 rpm at 4 °C for 30 min to collect plasma. Steroids were extracted from 10 μl of serum with the Steroid Displacement Reagent (Enzo Life Sciences) and corticosterone levels were measured by an enzyme immunoassay, according to the manufacturer’s protocol (Corticosterone Kit; Enzo). The minimal detectable concentration of corticosterone was 27 pg/ml. Optical densities were read at 405nm. The values were then calculated using a curve fit, ranging from 32 to 20000 pg/ml.

### Immunohistochemistry

Dams were perfused transcardially with saline (0.9%), followed by buffered paraformaldehyde solution (4%, pH 7.4). Brains were post-fixed for 4 h and then incubated in 30% sucrose-PBS solution overnight. Thirty micrometer thick coronal sections were cut with a cryostat (Leica Biosystems) and stored in anti-freeze solution (PBSx2 + 30% ethylene glycol + 20% glycerol) at -20°C until use.

Immunolabeling for Cox-2 and Lipocalin-2 was performed on free-floating sections using the avidin-biotin-HRP system with 3, 3’-diaminobenzidine tetrahydrochloride (DAB; Sigma-Aldrich) as chromogen, according to standard protocols. Briefly, the sections were first incubated in either rabbit anti-Cox-2 (R1747, 1:1000; Santa Cruz Biotechnology) or goat anti-Lipocalin-2 (MAB1757; 1:1000; R&D systems) overnight at room temperature, followed by biotinylated goat anti-rabbit IgG (1:1000; Vector Labs) or horse anti-goat IgG (1:1000; Vector Labs), for 2 h at room temperature. Sections were then incubated in avidin-biotin complexes (1:1000; Vector Labs) for 2 h followed by 0.5mg/ml DAB, hydrogen peroxide (0.01%) and nickel ammonium sulfate (2.25%) in sodium acetate buffer (0.1 M, pH 6.0) for 6 min.

Sections were analyzed on a Nikon 80i microscope equipped with epifluorescence, using 10X or 20X objectives. Quantification of cyclooxygenase-2 (Cox-2) and lipocalin immunoreactivity was performed in two anatomical sites (hypothalamic and the striatal area) of both hemispheres. The number of Cox-2 and lipocalin positive cells was counted in full 10X fields and averaged for each mouse. Captured images were processed in Adobe Photoshop (Adobe Systems) with adjustments of brightness and contrast.

### Statistical analysis

Data analysis was carried out in Graphpad Prism v. 6.01. Results are presented as mean ± SEM. Statistical comparison of two groups was done using Mann–Whitney test and for more than two groups Kruskal-Wallis 1-way ANOVA.. When having two categorical variables, 2-way ANOVA was performed, followed by Šídák post-hoc analysis multiple comparisons tests. Pearson correlation coefficients were used to examine the relationship between inflammatory gene expression in the hypothalamus and corticosterone levels in dams exposed to sick pups. *P* < 0.05 was considered statistically significant.

## Results and discussion

To investigate if inflammation, or inflammatory responsiveness, can be socially transmitted we used mice injected with LPS as a model. First, we focused on how dams react to sick offspring, due to their tight bond and the importance of their interaction for the survival of the pups. We induced systemic inflammation in 8-9 days old mouse pups by intraperitoneal injection of LPS (Fig. 1A). This procedure increased pup-directed support behaviors (Fig. 1B-D and Suppl. Fig. 1) and corticosterone levels (Fig. 1E) in the dams, showing that the dams detected and responded to the challenged state of the pups. After 3h of interaction with the immune challenged pups, the levels of inflammatory transcripts in the hypothalamus of the dams were examined. Surprisingly, we found a strong induction of inflammatory genes such as *Ptgs2* (encoding cyclooxygenase-2), *Il1b, Tnf, Il6, Ccl2* and *Cxcl10* (Fig. 1F). Furthermore, we detected a strong induction of PTGS2/cyclooxygenase-2 and Lipocalin-2 protein in brain endothelial cells (Fig. 1G-L). In several replicates of the experiment, we observed that the inflammatory response induced by sick pups and the fraction of responsive dams varied within and between experiments (Fig. 1M-O). We were unable to identify the factors underlying this variability, but we never detected an inflammatory state in a dam with saline-injected pups, indicating that the effect was specific to dams with sick pups. Induction of cytokine and chemokine mRNAs was also detected in the liver (Fig. 1P) and other brain regions (hippocampus and nucleus accumbens; Suppl. Fig. 2) of dams exposed to sick pups.

**Fig. 1.**
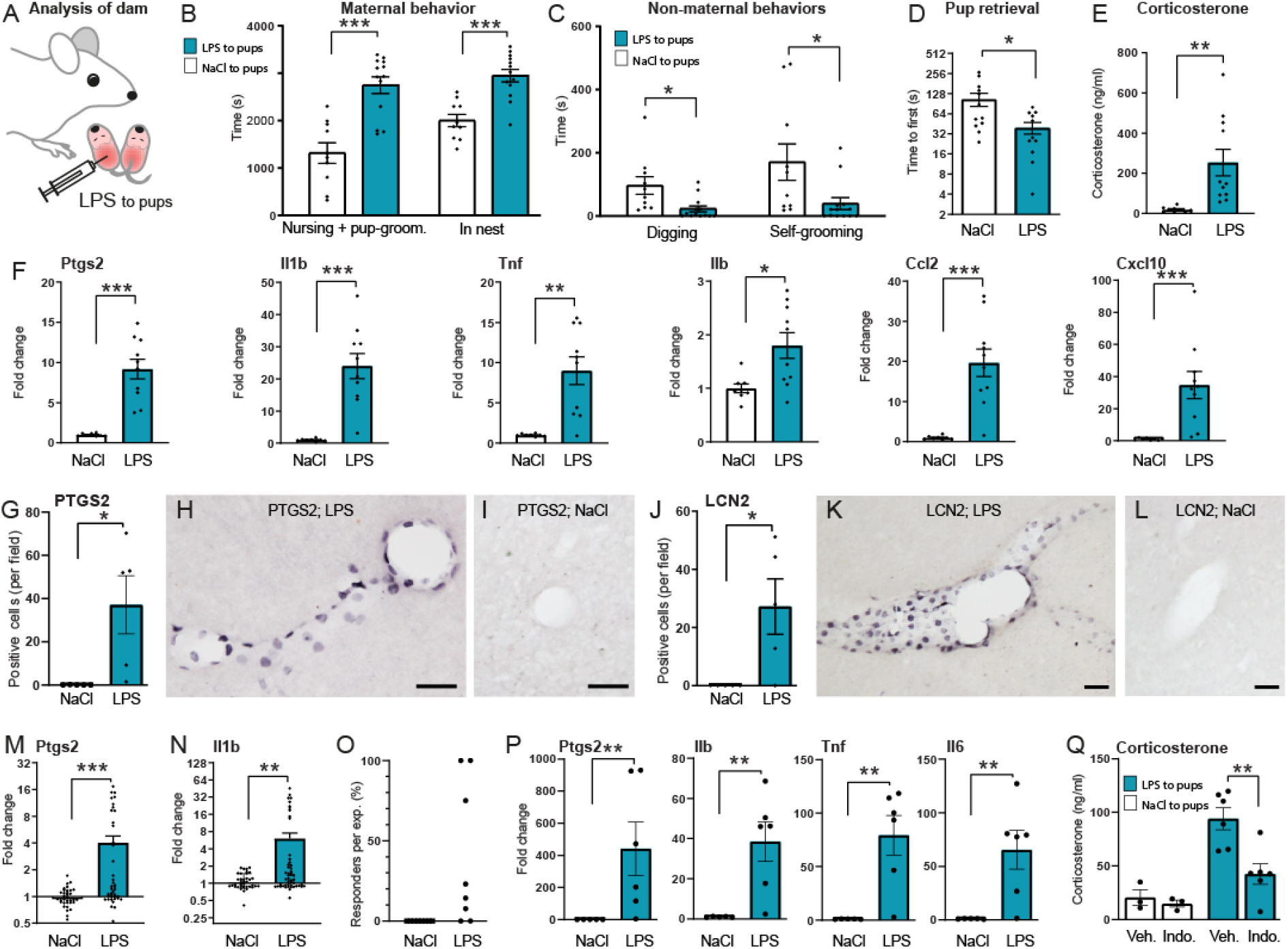
Exposure to immune challenged pups elicited a central and systemic inflammatory response in dams. Dams exposed to immune challenged pups (A) displayed increased maternal-related behaviors at the expense of other behaviors (B-D; n = 8-13/group), and showed increased plasma levels of corticosterone (E; n = 9-10/group)). They also displayed an induction of cytokine and chemokine mRNA in the hypothalamus (F; n=8-10/group) and an induction of PTGS2 and Lipocalin-2 protein in the brain endothelium (G-L; quantification of profiles in G, J, n=6/group; and representative pictures from the immunohistochemistry in H, I, K, L). The induction of inflammatory genes in the hypothalamus varied between individuals (M, N; n = 34-41 per group) and experimental batches (O). A strong induction of cytokines and chemokine mRNA was also found in the liver of dams exposed to sick pups (P; n=5-6/group). The corticosterone release induced by exposure to sick pups was attenuated by the cyclooxygenase inhibitor indomethacin (Q). Indo.; indomethacin, Lcn2; lipocalin-2, LPS; Lipopolysaccharide, NaCl; Saline, Veh; vehicle. Scale bar = 50 μm. Data are expressed as mean ± SEM. Gene expression changes are expressed as fold change versus the control group. Significance was determined with Mann-Whitney test, or Two-way ANOVA followed by Sidak posthoc test, with \**P* < 0.05, \*\**P* < 0.01, \*\*\**P* < 0.001.

**Fig. 2.**
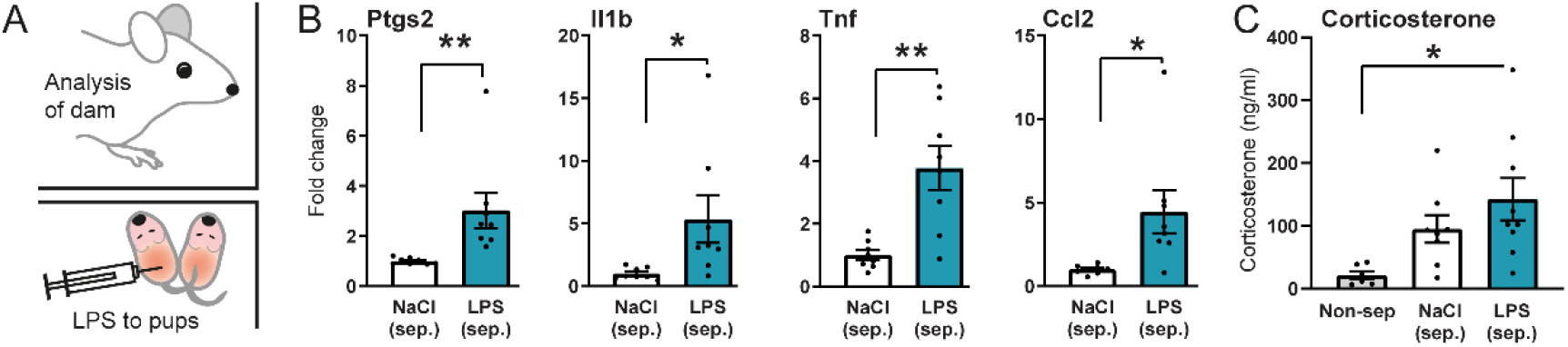
Social transmission of an inflammatory response does not require physical contact between dams and pups. The induction of cytokine and chemokine mRNA in the hypothalamus could also be observed when pups were injected with LPS and kept in a separate cage next to the cage with the dam (A, B n=8/group). Corticosterone levels were increased in dams with separated pups subjected to immune challenge (C; n = 6-8/group). LPS; Lipopolysaccharide, NaCl; Saline. Gene expression changes are expressed as fold change versus the control group. Data are expressed as mean ± SEM. Significance was determined using Mann-Whitney test for (B) and One-Way ANOVA (C) followed by Sidak posthoc test if applicable with *P < 0.05, **P < 0.01, *** P < 0.001.

We next investigated the relation between the transmitted inflammatory response and the elevation of corticosterone in the dams with immune challenged pups. Corticosterone levels strongly correlated with cytokine levels in the hypothalamus (Suppl. Fig. 3). To get an indication of whether the transferred inflammation mediated the stress hormone release in the dams exposed to immune challenged pups (in the same cage), we inhibited inflammatory signaling in the dams with the prostaglandin synthesis inhibitor indomethacin (Fig. 1Q), which previously has been shown to inhibit inflammation-induced corticosterone release (9). Indeed, indomethacin attenuated the increase in corticosterone levels induced by exposure to immune challenged pups (Fig. 3B) indicating a causal role of the inflammatory signaling for the stress hormone response.

**Fig. 3.**
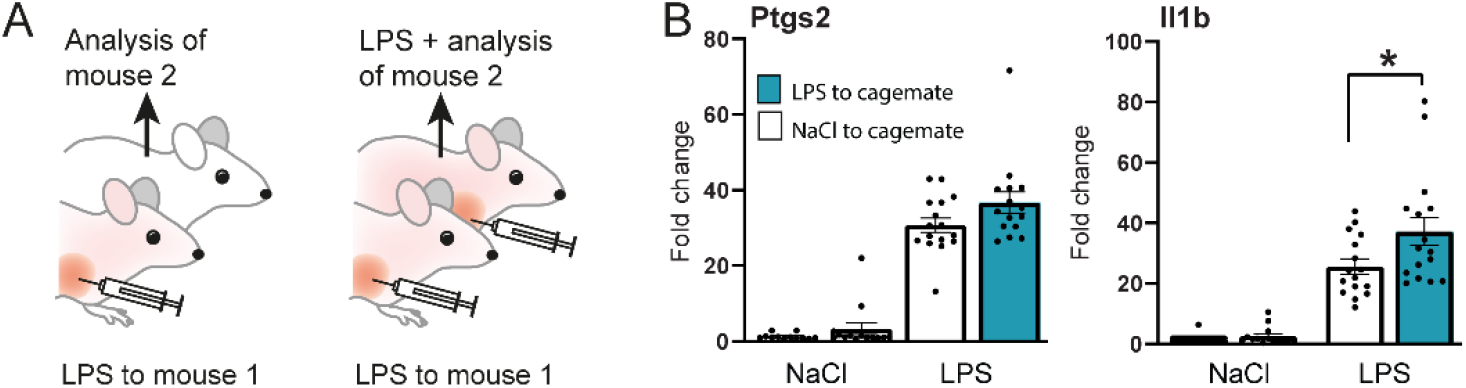
Interaction with an adult immune challenged conspecific induced an increased responsiveness to immune challenge. Exposure to an adult sick cage-mate (A) potentiated aspects of the inflammatory response to LPS (B; n=13-16/group). LPS; Lipopolysaccharide, NaCl; Saline, Veh; Vehicle. Data are expressed as mean ± SEM. Significance was determined using Two-way ANOVA followed by Sidak posthoc test with *P < 0.05

To investigate if the inflammatory signaling in the dams required physical contact with the pups, we moved the pups to a separate cage next to the cage with the dam, injected them, and kept them there for the entire experiment. Despite the lack of physical contact between pups and dam, we found an induction of inflammatory genes in the hypothalamus of the dams (Fig. 2B, C). Thus, the transferred inflammatory signaling was not due to LPS excretion from the pups, or any mechanism involving direct physical contact between pups and dam. We found no induction of inflammatory genes in dams with separated pups given saline (Fig. 2B), suggesting that stress induced by the separation is not sufficient for such an induction. Although we did not identify the route of transmission, olfactory cues are likely candidates since they have been shown to be important for avoidance of sick animals (10) and social transfer of stress and hyperalgesia (1-3). In the latter case, the signal was shown to spread between cages (2), similar to what we found in the present study. However, we cannot exclude other routes of transmission such as visual cues (4,11) or vocalizations (12).

Next, we investigated if a similar transfer of inflammation takes place between adult mice (Fig. 3A). In dyads of females, we injected one of the mice with LPS to induce sickness, and assessed the cytokine mRNA levels in the hypothalamus of the cage-mate. We observed no consistent induction of inflammatory genes in the bystander mice (Fig. 3B) indicating that the social transfer of inflammatory signaling, as observed between pups and dams, is not present in all ages and kinships. However, when bystander mice were immune challenged after exposure to a sick cage-mate, we observed a mildly potentiated induction of IL-1β mRNA in the hypothalamus (Fig. 3B). Similar potentiation of inflammatory signaling have been observed previously in animals interacting with sick conspecifics (13). The potentiation in adults can be seen as a “less strong” form transmission and is conceptually similar to “transmitted pain”, in which a mouse in pain induces increased sensitivity to pain (1, 2), but not spontaneous pain, in bystanders. It is unclear why the “strong form” of transmission, i.e. induction of inflammatory signaling in an unchallenged bystander, took place between pups and dams but not between adult females. Perhaps it reflects an adaptation in the dam to the necessity of close physical contact with sick pups, or a mechanism for supporting them by mediators through the milk (14).

For stress, fear and pain, the social transfer has been explained in terms of an adoption of the sensory-affective state of a partner in distress. Such mechanisms could also be at play for the transfer of inflammatory signaling that we observe here. However, this inflammatory mirroring is different from the other examples of transfer in the sense that it not only involves activity in the neurons underpinning sensory-affective states but also a strong and broad induction of inflammatory mediators in the brain and periphery. Such induction is unlikely to be a generic stress effect since separation of the dams from the pups, which is stressful for the dams, was not enough to trigger any inflammatory response. Furthermore, although some strong and persistent stress paradigms can induce low-grade inflammatory signaling (15,16), the magnitudes of such immune activations are much lower than the ones we observe here.

We conclude that dams exposed to immune challenged pups respond with a strong systemic induction of inflammatory genes. The induction contributed to the stress hormone response in the dam and did not rely on physical contact between dams and pups. These observations show that systemic inflammation can be socially transmitted. We believe our findings will open new avenues for research on social behavior, just like the identification of similar concepts such as observational fear and transmitted pain has done.

## Acknowledgments

This study was supported by the Knut and Alice Wallenberg foundation, the Swedish Research Council (2022-00952 and 2020-00881), the Swedish Brain Foundation (FO2022-0114 and FO2021-0010), and Lions forskningsfond mot folksjukdomar. We thank Eda Erdem, Sofie Arvidsson and Ellen Thunholm for their assistance.

## Notes

### Competing Interest Statement

The authors have declared no competing interest.

