## Supplemental data for "Social transmission of inflammation in mice"

### Supplementary data

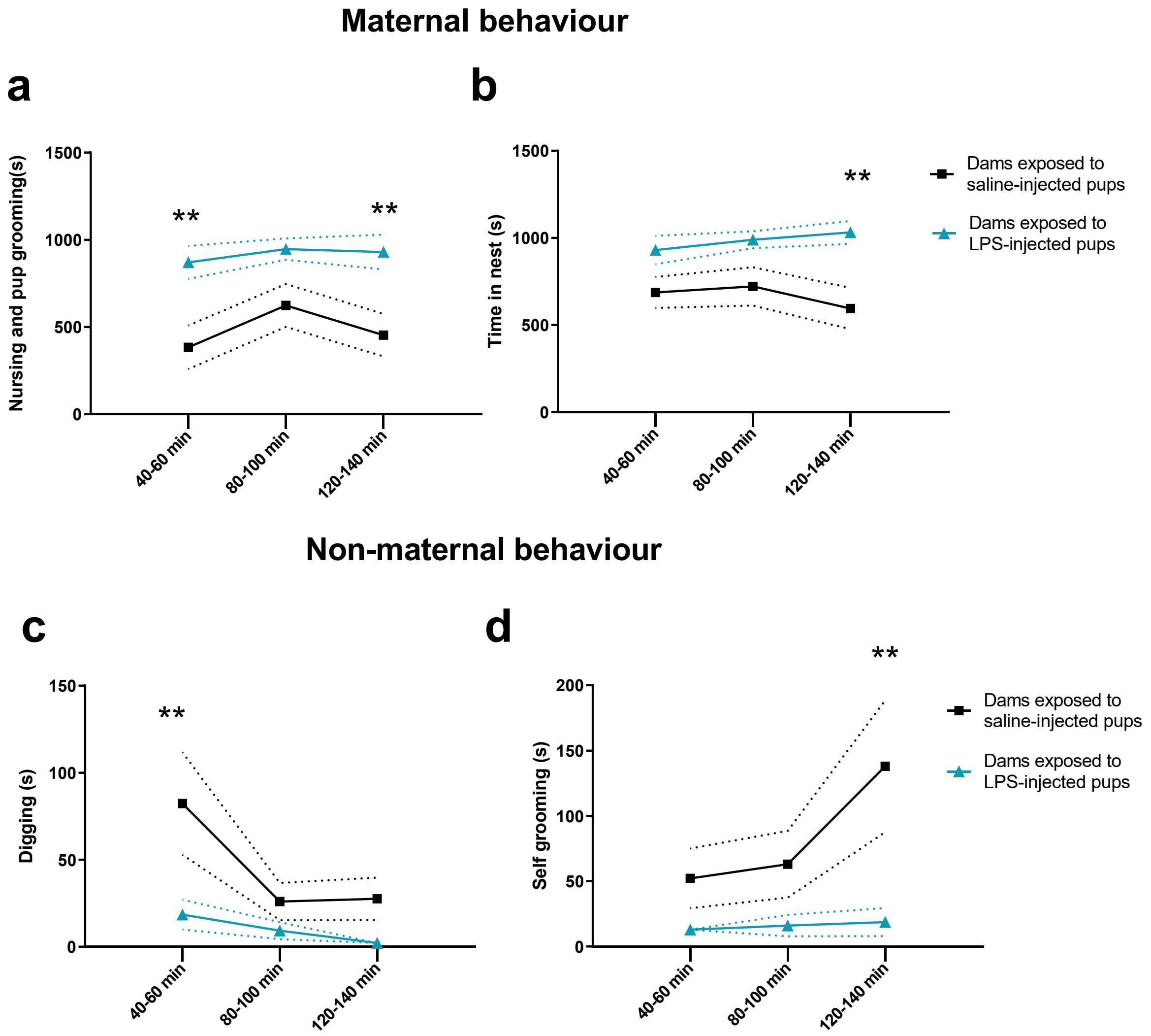


**Supplementary Fig. 1. Time course of maternal and non-maternal related behaviors of dams exposed to immune challenged pups**. Time course of a) nursing and pup grooming behavior, time in the nest, c) digging and d) self-grooming displayed by dams during the 3h exposition to LPS (N dams=13) or saline injected pups (N dams=10) in time bins of 20 min. c) Time course of digging and d) self-grooming performed by dams exposed to LPS-injected pups. Two-way ANOVA followed by Šidák posthoc test was performed. ** $p < 0{,}01$.

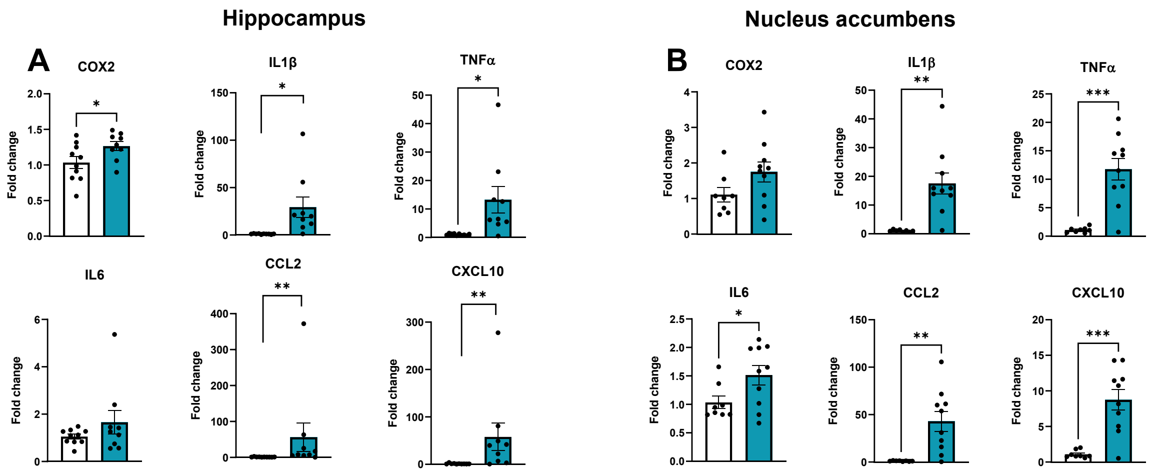


**Supplementary Fig. 2. Dams exposed to immune callenged pups show a strong induction of inflammatory genes in the hippocampus and nucleus accumbens.** (a) Hippocampal expression of Cox2, IL1β, TNFα, CCL2, CxCL10 was significantly increased in dams exposed to LPS injected pups. (b) In the nucleus accumbens (B), significant increases in the expression of IL1β, TNFα, IL6, CCL2, CxCL10 were found in dams exposed to sick pups. Mann-Whitney test. *$p < 0.05$, **$p < 0.01$, ***$p < 0.001$

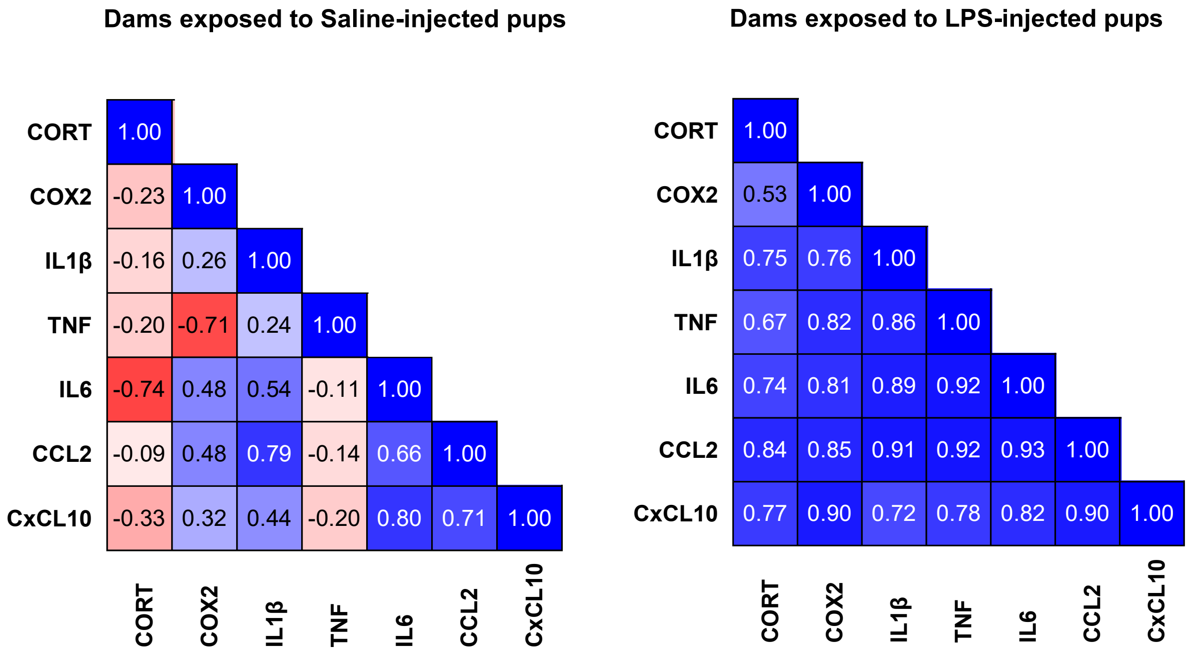


**Supplementary Fig.3. Corticosterone levels and hypothalamic inflammatory gene transcripts strongly correlate in dams exposed to LPS-injected pups.** Pearson correlation matrix was calculated. A decision coefficient (r2) was determined by the Pearson correlation coefficient.
